# Nicotine interacts with DNA lesions induced by alpha radiation which may contribute to erroneous repair in human lung epithelial cells

**DOI:** 10.1101/2024.09.07.611801

**Authors:** Nadia Boroumand, Carol Baghdissar, Karine Elihn, Lovisa Lundholm

## Abstract

**Purpose:** Epidemiological studies show that radon and cigarette smoke interact in inducing lung cancer, but the contribution of nicotine in response to alpha radiation emitted by radon is not well understood.

**Materials and methods:** Bronchial epithelial BEAS-2B cells were either pre-treated with 2 μM nicotine during 16 h, exposed to radiation, or the combination. DNA damage, cellular and chromosomal alterations, oxidative stress as well as inflammatory responses were assessed to investigate the role of nicotine in modulating responses.

**Results:** Less γH2AX foci were detected at 1 h after alpha radiation exposure (1–2 Gy) in the combination group versus alpha radiation alone, whereas nicotine alone had no effect. Comet assay showed less DNA breaks already just after combined exposure, supported by reduced p-ATM, p-DNA-PK, p-p53 and RAD51 at 1 h, compared to alpha radiation alone. Yet the frequency of translocations was higher in the combination group at 27 h after irradiation. Although nicotine did not alter G2 arrest at 24 h, it assisted in cell cycle progression at 48 h post radiation. A slightly faster recovery was indicated in the combination group based on cell viability kinetics and viable cell counts, and significantly using colony formation assay. Pan-histone acetyl transferase inhibition using PU139 blocked the reduction in p-p53 and γH2AX activation, suggesting a role for nicotine-induced histone acetylation in enabling rapid DNA repair. Nicotine had a modest effect on reactive oxygen species induction, but tended to increase alpha particle-induced pro-inflammatory IL-6 and IL-1β (4 Gy). Interestingly, nicotine did not alter gamma radiation-induced γH2AX foci.

**Conclusions:** This study provides evidence that nicotine modulates alpha-radiation response by causing a faster but more error-prone repair, as well as rapid recovery, which may allow expansion of cells with genomic instabilities. These results hold implications for estimating radiation risk among nicotine users.

## 1. Introduction

Human lungs are consistently subjected to inhaled environmental and chemical substances that potentially could damage the DNA in cells and/or promote pre-existing damage (Hill et al., 2023; Shankar et al., 2019). Radon, the second largest cause of lung cancer after smoking (Darby et al., 2005; Vogeltanz-Holm and Schwartz, 2018), finds its place among the top environmental hazards to public health and has been considered a human carcinogen by the World Health Organization (WHO) since 1987 (Vogeltanz-Holm and Schwartz, 2018; WHO, 2009). Approximately 3–14 % of lung cancer cases are caused by radon (WHO, 2009).

The solid products of radon decay tend to accumulate on the bronchial wall, the primary site for lung cancer development (Hofmann et al., 1990; Truta-Popa et al., 2011; Madas, 2016). Alpha-ionizing radiation emitted by radon and its decay products has been associated with a wide array of cytotoxic and genotoxic effects. The energy deposited by these alpha particles (He^2+^ ions) is sufficient to penetrate the bronchial epithelia and deliver high local doses to the cells and DNA, even though the average lung dose might be considered to be low (Hofmann et al., 1990; Truta-Popa et al., 2011; Madas, 2016). Direct ionizations caused by alpha particles are more prone to cause clustered DNA damage compared to damage induced by gamma or X-rays (National Research Council US Committee on Health, 1999). Complexity involved in repairing clustered damages can play a role in the onset of cancer (Durante, 2009). Due to the deleterious effects of ated alpha particles in the bronchi, WHO has defined a national annual average residential radon concentration reference level of 100 Bq/m^3^, with stating that if this limit is not applicable under the prevailing country-specific conditions, it should not exceed 300 Bq/m^3^ (World Health Organisation, 2009). However, it is known that other stressors such as smoking can increase the lung cancer risk induced by radon, which can lead to inaccuracies in the derived risk estimates (Nilsson and Tong, 2020; Kreuzer et al., 2018).

In addition to the specific risks posed by radon exposure, epidemiological studies have shown that cigarette smoke (CS) amplifies the risk of radon-induced lung cancer both regarding occupational (Leuraud et al., 2011) and residential exposure (Darby et al., 2005). Statistically, the joint effect of these two agents appears to be best described by a less than multiplicative interaction type (Council, 1999; Su et al., 2022). Moreover, with respect to sex-related differences, studies have shown that smoking can increase the risk of developing lung cancer 22-fold in men and 12-fold in women (George and Shoenfeld, 2000). A positive correlation was reported between androgens and incidence of lung cancer in older men (Chan et al., 2017). These findings along with many others (Chan et al., 2017; Fucic et al., 2010; Chang et al., 2014) indicate a possible role for hormonal status in the etiology of lung cancer. More advanced-stage adenocarcinomas were seen in pre-menopausal women compared to post-menopausal women and men, also supporting a role of estrogens (Rodriguez-Lara and Avila-Costa, 2021).

CS-induced cancer is driven by more than 60 carcinogens that directly cause damage and mutations in the DNA. These include tar, arsenic, 1,3-butadiene, carbon monoxide, nitrosamines, aldehydes and small organics (Tweed et al., 2012). However, nicotine has often been regarded as a comparatively safer alternative to CS, which has provoked a debate within the medical community as to giving rise to consumption of nicotine through various alternative products, including E-cigarettes and smokeless tobacco such as snus. There has been a rapid increase in nicotine vaping among the younger population during recent years, 11 % of US high school students were e-cigarette users in a 2021 survey (Gentzke et al., 2022). Consequently, growing numbers of studies are examining its possible correlation with cancer incidence (Grando, 2014).

Nicotine is the primary addictive compound that contributes to persistent tobacco use, through inducing release of catecholamines into the bloodstream leading to improved physical and mental ability (Mohamed et al., 2023; Abdel-Rahman Mohamed et al., 2022). Simultaneously, it can provide a tumor-supporting microenvironment favoring cell proliferation, survival and metastasis, all of which can provoke aberrant signaling and support expansion of cells with genomic instabilities (reviewed in (Chernyavsky et al., 2015)). Studies have shown that several tobacco-specific nitrosamines derived from nicotine are carcinogenic such as NNK [4- (methylnitrosamino)-1-(3-pyridyl)-1-butanone] and NNN (N′-nitrosonornicotine) (Pérez-Ortuño et al., 2016). Although nicotine itself is not considered to initiate cancer, studies suggest that both nicotine and NNK can induce DNA adducts such as γ-OH-PdG and O6-medG which are often repaired by NER and BER repair mechanisms (Lee et al., 2018). This can lead to collapse of replication forks upon collision with these primary DNA lesions. One of the possible ways that nicotine can induce the formation of these adducts is through redox mechanisms by oxidizing DNA bases (Ginzkey et al., 2012). Besides genotoxic mechanisms (Tweed et al., 2012; Grando, 2014), impairing the DNA-damage response (DDR) may be another mechanism through which nicotine exerts its effect (Lee et al., 2018; Tang et al., 2022). Furthermore, nicotine exposure has been indicated to exert pro-survival effects on lung epithelial cells through the activation of intracellular growth factors such as protein kinase C (PKC), or by increasing the activity of Bcl-2 to counteract apoptotic signals (Mai et al., 2003). Nicotine was also shown to block the suppression of PKC and extracellular signal-regulated kinase (ERK2) activity caused by chemotherapy, having adverse effects on the outcome of the treatment (Mai et al., 2003). In parallel with this data, multiple studies have provided evidence that continuing smoking following a cancer diagnosis can adversely impact the overall survival, tumor control, treatment side effects and occurrence of secondary cancers among patients undergoing radiotherapy, showing a possible interaction also between smoking and X-ray radiation (Perdyan and Jassem, 2022; Wong et al., 2020). The proven role of nicotine, a major component of cigarettes, in altering cellular functions in favor of cancer development strongly supports the candidacy of this compound as a potential target for study.

Here we used BEAS-2B cells, an established *in vitro* lung model (Lee et al., 2018; Park et al., 2015) having functional p53 signaling (Park et al., 2015). We showed that nicotine promoted an enhanced proliferative potential compared to alpha-irradiated cells without nicotine treatment. To investigate this difference, we examined if the DNA damage/repair dynamics differed between the aforementioned groups. The assessment demonstrated a reduced activation/expression of DDR proteins in the combination group. This reduced response was associated with less fragmented DNA and a higher translocation rate, indicating a faster but error-prone repair. Interestingly, inhibiting histone acetylation revealed that chromatin structure is a modifying factor in the nicotine-accelerated repair function. Collectively, our data indicate that nicotine enables cells to recover faster after alpha-particle induced DNA damage at the cost of erroneous repair and this might be mediated through a more relaxed chromatin structure. This investigation can shed light on the combined effects of nicotine and radiation, offering valuable insights into the mechanisms and risks associated with their interaction.

## 2. Materials and methods

### 2.1. Cell culture and chemical treatment

Human normal lung bronchial epithelial BEAS-2B cells, immortalized by adenovirus 12-SV40 (kindly donated by assoc. prof Hanna Karlsson, Institute of Environmental Medicine, Karolinska Institutet, Sweden), were cultured in airway epithelial cell growth medium with supplement mix and 1 % penicillin-streptomycin (10,000 U penicillin and 10 mg streptomycin/ml) (all from Sigma-Aldrich, Schnelldorf, Germany). Cells were grown at 37 °C and 5 % CO_2_ and were passaged every two days in 75 cm^2^ culture flasks at a seeding density of 8.0 × 10^5^ cells. BEAS-2B cells reached 60–70 % confluence before the start of each experiment. One day before irradiation, 2.0 × 10^5^ cells for γH2AX assay and 3.5 × 10^5^ cells for the rest of the assays were seeded on squared (22 ×22 mm), and rounded (d=32 mm) glass coverslips, respectively. As for the nicotine group and combination group, 2 μM nicotine treatment (Sigma-Aldrich) was started 16 h before irradiation. Cells were continuously receiving nicotine treatment after irradiation in all experiments except those in Fig. 1D, E, H, 2A, 6A, 4D and S1. This alteration in methodology arose during the study and was integrated subsequently, after several experiments had already been conducted. Therefore, it became impractical to repeat or modify the earlier sets of experiments with the evolved protocol.

**Fig. 1.**
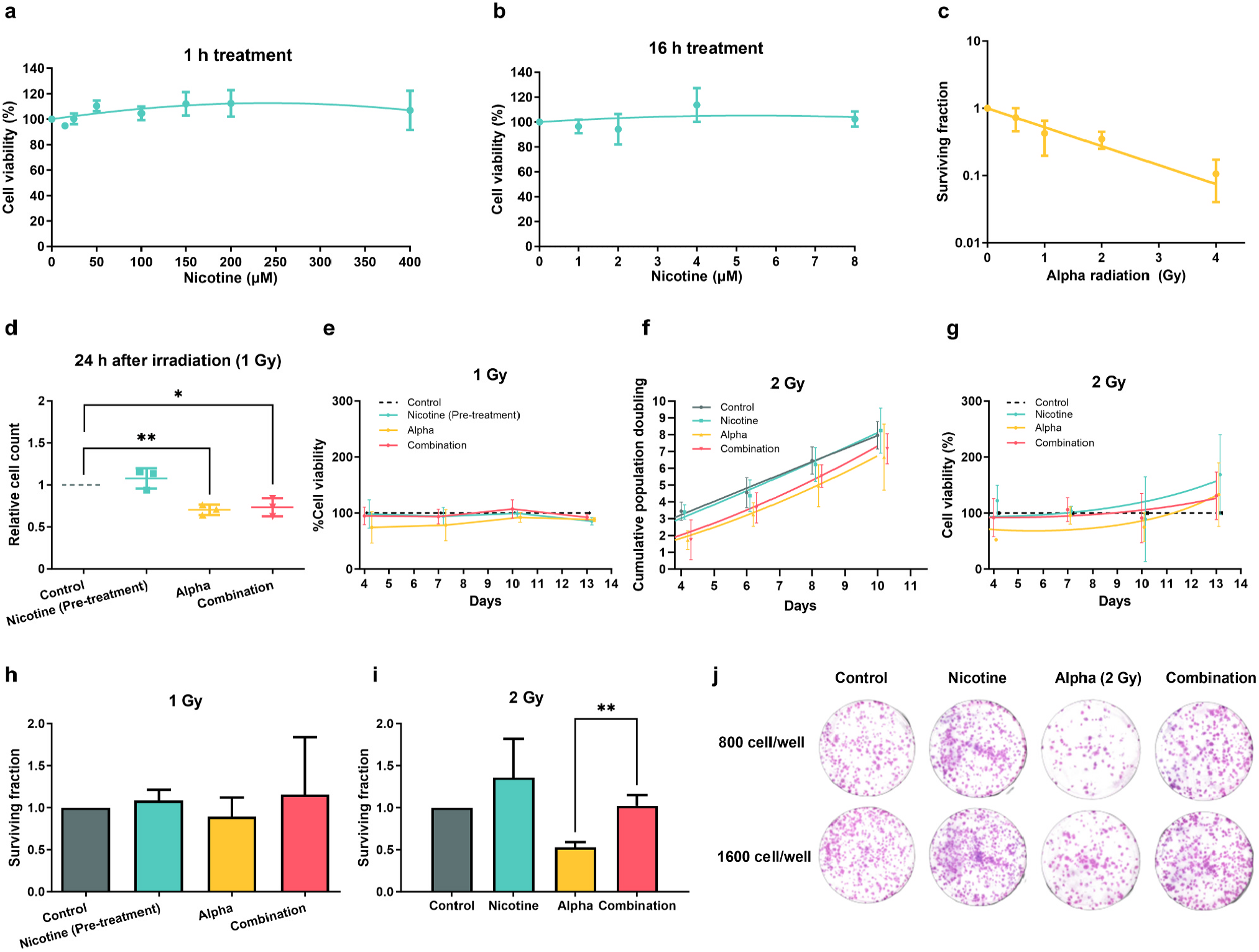
Nicotine promotes faster recovery and enhanced survival in alpha particle-irradiated cells. a, b) Toxicity of nicotine in BEAS-2B cells treated with 15–800 μM for 1 h (a), and 1–8 μM for 16 h (b). c) Clonogenic survival of BEAS-2B cells 11 days post irradiation with 0, 0.5, 1, 2, and 4 Gy doses of alpha particles. d) Trypan blue exclusion cell counts 24 h post irradiation in cells pre-treated with 2 μM nicotine for 16 h, alpha particles (1 Gy), or in combination. e) Cell viability during 13 days post irradiation in cells pre-treated with 2 μM nicotine for 16 h, alpha particles (1 Gy), or in combination. f, g) Cumulative population doubling and cell viability in cells treated with 2 μM nicotine (pre- and post-treatment), alpha particles (2 Gy), or in combination during 10 and 13 days post irradiation, respectively. h, i) Colony-forming capacity of cells treated with 2 μM nicotine (pre-treatment for 1 Gy, pre- and post-treatment for 2 Gy), alpha-irradiated (1, 2 Gy), or in combination, seeded at concentrations of 400, 800, 1600 cells per plate, and kept for 11 days. j) Representative images of colonies grown during 11 days after the respective treatments. Data are expressed as means ± SD from at least three independent experiments. The significance level was p < 0.05 (*) and p < 0.01 (**) (two-tailed Student’s t-test).

The hormone treatment was administered at a final concentration of 10 nM 17β-estradiol (E2) (E8875, Sigma-Aldrich) or 10 nM 5α-dihydrotestosterone (DHT) (NMID680, Sigma-Aldrich), which were added after irradiation until further processes. Doses that are sufficient to activate receptors and act via hormone response elements were selected based on previous literature (La Sala et al., 2010; Chhipa et al., 2009). Cell viability and survival were evaluated at 10 and 11 days post irradiation, respectively.

PU139, a pan-histone acetyltransferase (HAT) inhibitor (2-(4-Fluorophenyl)isothiazolo[5,4-*b*]pyridin-3(2 H)-one, HY-124696, MedChemExpress) treatment was performed for the respective endpoints using 5 μM PU139 for 16 h before irradiation. The drug was kept in the media thereafter until 1 h after irradiation when the cell pellets were collected.

### 2.2. Radiation exposure

Irradiation of cells was performed using an alpha source (^241^Am source, 50.0 ± 7.5 MBq, Eckert & Ziegler Isotope Products GmbH, Germany) with a dose rate of 0.223 Gy/min and an average linear energy transfer (LET) of 90.9 ± 8.5 keV/μm, as described earlier (Cheng et al., 2018; Staaf et al., 2012a). Additionally, in each experiment, a non-exposed control group and a nicotine-only treated group were included. After exposure, the coverslips with cells were transferred to new 6-well plates and were kept inside the incubator at 37 °C and 5 % CO_2_ until further processing.

Gamma radiation was delivered to the cells from a ^137^Cs source Gammacell® 40 Exactor (MDS Nordion, Ontario, Canada) at a dose rate of 0.70 Gy/min. Control samples underwent comparable treatment as the irradiated samples, excluding the actual irradiation.

### 2.3. Trypan blue exclusion assay and population doubling

The trypan blue exclusion assay was employed to determine the cell density and viability throughout the course of an experiment, using an automated cell counter (Countess, Invitrogen, Paisley, UK). Population doubling was calculated for cell growth monitoring using: *PD* = *ln (Nt/ N0) / ln2* where Nt is equal to the concentration of cells after the harvest, N0 is equal to concentration of cells seeded. Thereafter, cumulative increase in population doubling was accounted for, creating cell growth curves.

### 2.4. MTT assay

Cell viability was assessed using 3-(4,5-dimethylthiazolyl-2)-2,5-diphenyltetrazoliumbromide reagent (MTT) assay (Sigma-Aldrich). After the treatment of interest or sham treatment, cells were seeded in optical clear 96-well flat bottom plates. At the respective time point, MTT reagent was added at a final concentration of 0.25 mg/ml and incubated for 4 h at 37 °C. Thereafter, 10 % sodium dodecyl sulphate (SDS) in 0.01 M HCl was added to dissolve the purple formazan crystals. A spectrophotometric microtiter plate reader was utilized to measure the absorbance at 570 nm (BMG LABTECH, Germany). The data obtained from both control and treated cells were analyzed and presented as a percentage of viable cells.

### 2.5. Colony formation assay

Clonogenic survival assay was performed as described earlier (Olofsson et al., 2020). Briefly, cells were seeded at different cell densities ranging from 400 to 1600 cells per well in duplicates in 6-well plates. Following an 11-day period, the colonies were fixed and stained using 25 % methanol and 5 % Giemsa solution (Sigma-Aldrich). The countPHICS software was employed to score the number of colonies per well (Brzozowska et al., 2019).

### 2.6. γH2AX immunofluorescence assay

Following irradiation, cells were kept at 37 °C to allow repair. Cells were fixed after 30, 60, 120, 240, 360 min and 24 h using 70 % ethanol, after which immunostaining was carried out as described earlier (Staaf et al., 2012b). Briefly, cells were permeabilized in 2 % Triton X-100 (Sigma-Aldrich), incubated with anti-phospho-histone H2AX antibody Ser139 (16–202 A, Sigma-Aldrich), followed by secondary antibody (F0257, anti-mouse IgG conjugated to FITC, Sigma-Aldrich). Antibodies were diluted 1:800 (1.25 μg/ml) and 1:200 (5 μg/ml), respectively, in PBS containing 2 % bovine serum albumin (BSA) Fraction V (Sigma-Aldrich). To provide a counterstain for cellular nuclei, coverslips were mounted using VECTASHIELD® Mounting Medium supplemented with 4’,6-diamidino-2-phenylindole (DAPI) (Vector laboratories, US).

### 2.7. Image acquisition and analysis of γH2AX foci

For each experiment, individual cells were randomly selected and captured using a Nikon Eclipse E800 fluorescence microscope equipped with a 100 × oil immersion lens (Nikon Eclipse E800, Nikon, Tokyo, Japan) and the ISIS image analysis system (Metasystems, Altlussheim, Germany) connected to a CCD camera. To analyze the images, a modified macro designed for ImageJ software version 1.43 u (Markova et al., 2007) was employed. The macro was used to determine the area and count the number of γH2AX foci. The foci were classified into two categories based on their areas, small (0.5–59 pixels) and large (60–500 pixels), as described earlier (Sollazzo et al., 2017). In each experiment, at least 100 cells were analyzed per condition.

### 2.8. RNA isolation and real time PCR analysis

RNA was extracted using the E.Z.N.A. Total RNA Kit I (Omega Biotek, Norcross, GA, USA). cDNA was synthesized using the High-Capacity cDNA Reverse Transcription Kit (Thermo Fisher Scientific) with random hexamer primers. Duplicate reactions of primers, cDNA and 5 × HOT FIREPol® EvaGreen® qPCR Supermix (Solis BioDyne) were mixed and ran on a LightCycler® 480 for real time PCR. The cycling was performed as: 95 °C (15 min), 40 cycles of 95 °C (15 s), 60 C (20 s) and 72 °C (20 s). The 2^-ΔΔCt^ method was used for the calculation of relative expression. The housekeeping 18S gene was employed for normalization and the specificity of the primers were tested by melting curve analysis. Forward and reverse sequences of primers are given earlier for 18S (Lundholm et al., 2014), IL-6 and IL-1β (Fromell et al., 2023) and CDKN1A (Lopez-Riego et al., 2023).

### 2.9. Mitochondrial ROS assay

Oxidative stress was quantified by using the dihydrorhodamine 123 (DHR123) assay as described earlier (Balaiya and Chalam, 2014). Briefly, cells were seeded at the density of 1000 cells/well and treated with 2 μM nicotine for 16 h. Following that, 100 μl freshly prepared working solution of DHR123 was added to each well. The cells were incubated for a total of 90 minutes and the fluorescence of Rh123 was measured after 20 minutes of incubation and every 10 minutes afterwards in order to determine the peak generation of reactive nitrogen oxide species (RNOS)/ROS at 485/528 nm (excitation/emission).

### 2.10. Alkaline comet assay

The alkaline comet assay was performed by mixing 1.5 × 10^5^ cells 1:1 with 2 % low melting point agarose (2-Hydroxyethylagarose, Type VII, Sigma-Aldrich, A4018) at 37°C, 100 μl of this suspension was placed on a microscope slide pre-coated with 0.5 % high melting point agarose (Type I-A, Sigma-Aldrich, A0169), covered with a coverslip and put at 4°C to solidify for 5 min. Thereafter the coverslips were removed and slides were immersed in a cold lysis buffer (2.5 M NaCl, 100 mM Na_2_EDTA, 10 mM Tris, 1 % Triton X-100, pH 10.0) for 1 h at 4 °C, followed by a brief rinsing in cold distilled water. Next, slides were randomly positioned in an EC 340 Maxicell Primo horizontal electrophoresis tank (Thermo EC, Holbrook, USA) filled with fresh, cold electrophoretic buffer (1 mM Na_2_EDTA and 300 mM NaOH, pH 13.3) and kept for 60 min for DNA unwinding. After that, electrophoresis was performed at 32.5 V (1 V/cm), 430 mA for 25 minutes. The number of slides in the unit was constant for all experiments. Slides were washed with neutralization buffer (0.4 M Tris, pH 7.5) and stained with DAPI. Comets were scored blindly using a 40 × objective on a Nikon Eclipse E800 fluorescence microscope. Images were analyzed using the Comet assay II software (Perspective Instruments, Suffolk, UK, version 2.11). At least 50 cells per experiment and time point were analyzed. The tail intensity (TI), also called % tail DNA, was selected for quantification of DNA damage. This measure describes the DNA fragment intensity in the comet tail versus the total intensity for that cell (head plus tail), expressed as percentage (OECD, 2016).

### 2.11. Western blot

After receiving the treatment of interest, cells on cover slips were incubated at 37 °C for 1 hour. Cells were then trypsinized, collected and lysed in RIPA buffer (50 mM Tris-HCl (pH 7.4), 150 mM NaCl, 0.5 % IGEPAL®, 5 mM EDTA (pH 8.0), 0.1 % SDS) supplemented with protease inhibitor cocktail, and PhosSTOP™ in case of assessing phosphorylation (Sigma-Aldrich, Germany). Proteins were separated using 4–12 % Bis-tris gradient gels in 1xMES buffer for smaller proteins and 3–8 % Tris-acetate gradient gels for larger proteins in NuPage® Tris-acetate running buffer (Invitrogen™, US) and were transferred to a nitrocellulose membrane (Thermo Scientific, US). The membrane was blocked using Odyssey® blocking buffer (LI-COR, Cambridge, UK) and Tris-buffered saline containing 0.05 % Tween (TBST), in 1:1 ratio, at room temperature for 1 h. Probing was performed with the following primary antibodies overnight at 4°C: pDNA-PK (pSer2056; SAB4504169, Sigma-Aldrich, 1:300), pATM (pSer1981; SAB4300100, Sigma-Aldrich, 1:500), p-p53 (pSer15; #9284, Cell Signaling Technology, 1:500), GAPDH (G8795, Sigma-Aldrich, 1:20 000), RAD51 (ab133534, Abcam, 1:10000), H3 (07–690, Sigma-Aldrich, 1:1000), H4 (07–108, Sigma-Aldrich, 1:1000), and modified histones including H3K9ac (#9677, Cell Signaling Technology, 1:1000), H3K9me3 (ab8898, Abcam, 1:1000) and H4K8ac (#2594, Cell Signaling Technology, 1:1000). Probing with secondary antibodies, infrared dye-conjugated goat anti-rabbit or donkey anti-mouse secondary antibodies (LI-COR, Cambridge, UK, 1:15000 diluted in TBST), was done for 1 h at room temperature. The membranes were scanned and protein levels were quantified with the Odyssey® S Infrared Imaging System (LI-COR) and analyzed with Image Studio™ Lite version 5.2 (LI-COR). The normalization was done by dividing levels of each protein to GAPDH. For histones, H3 and H4 were first normalized to GAPDH, then H3K9ac, H4K8ac and H3K9me3 were further normalized to the total histone proteins.

### 2.12. Flow cytometry

After the treatments of interest, cell cycle distribution was analyzed at 4, 24 and 48 h post exposure by Moxi GO II flow cytometer (MXG102, Orflo Technologies). Cells were harvested, fixed in cold 70 % ethanol and stored at - 20°C until further analysis. At the time of analysis, cells were resuspended in propidium iodide (PI) staining solution containing PBS with 20 μg/ml RNase (Sigma-Aldrich, Germany), 0.1 % Triton X-100 (Sigma-Aldrich, Germany) and 2 μg/ml PI (Sigma-Aldrich, Germany) and incubated at room temperature. Fluorescence was measured at 525/45 nm with a 561 nm low-pass filter. Data were analyzed using the software FCS Express (De Novo Software).

### 2.13. Chromosome preparation and fluorescence in situ hybridisation (FISH)

Alpha-irradiated and combination group cells were treated with 0.2 μg/ml colcemid during their last 3 hours in the incubator (out of totally 27 h). Cells were then harvested to prepare standard cytogenic slides. In brief, cells were resuspended in hypotonic solution (75 mM KCl) at 37 °C, fixed in 3:1 methanol:acetic acid, thereafter pelleted cells were diluted with fixative, and 300 μl of cell suspension was dropped on a clean microscope slide (Meher et al., 2023). The slides were subjected to the FISH assay according to the manufacturer’s protocol. For this assay, whole-chromosomepaint probes forchromosomes-1 and 2 (XCP 1 Orange, XCP 2 Green; XCyting chromosome Paints, MetaSystems GmbH, Germany) were utilized. The slides were scanned using a fluorescence microscope equipped with a 100x oil immersion lens (Nikon Eclipse E800). Image acquisition was performed with the auto exposure setting using a Cool Cube1 CCD camera and the ISIS image analysis system (MetaSystems GmbH). Translocations were analyzed 27 h after irradiation to allow cells to proceed through one mitosis. The frequency of translocations involving chromosome 1 and chromosome 2 with each other, or with other chromosomes stained with DAPI, was blindly assessed. Any reciprocal translocations were counted as two translocations. At least 100 metaphases were scored per slide.

### 2.14. Statistical analysis

Comparisons were conducted using unpaired t-test by GraphPad Prism ver. 9.3.1 (GraphPad Software, US), except for the distributions in Fig. 2C, 3B, and 6C, where a non-parametric Mann-Whitney test was used. A significance level of p < 0.05 was considered as statistically significant. MTT, cell growth and cell viability curves were fitted to a linear quadratic equation. The survival curves after alpha particle exposure were fitted to semilog line (X is linear, Y is log).

**Fig. 2.**
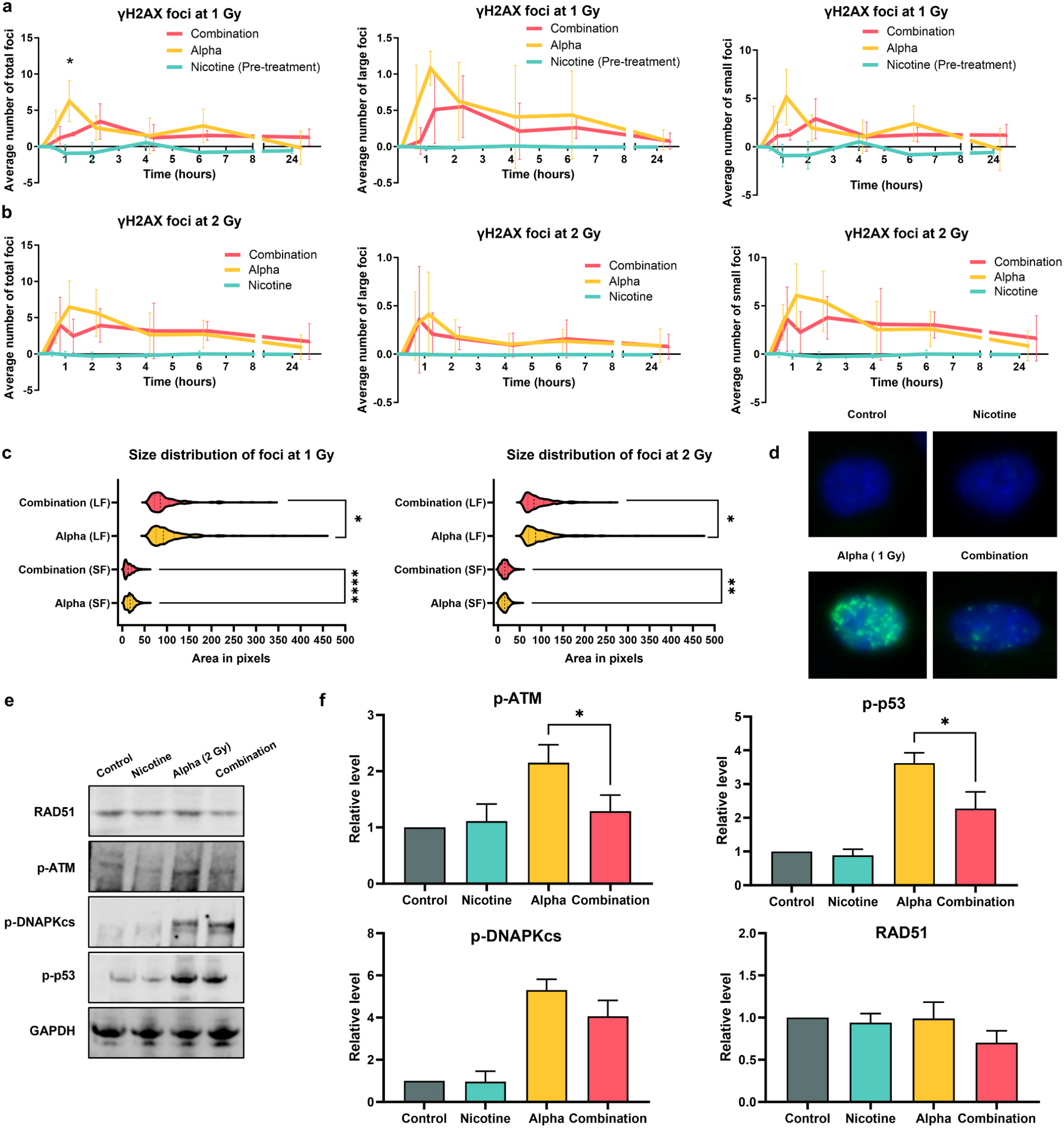
Nicotine alters the DNA damage/repair dynamics in alpha-irradiated cells. a, b) Kinetic response of γH2AX foci formation as a function of time over 24 h. Immunofluorescence was used to capture representative γH2AX focus images from control, 2 μM nicotine-treated cells (pre-treatment for 1 Gy, pre- and post-treatment for 2 Gy), alpha-irradiated cells (1, 2 Gy), and the combination. γH2AX foci were quantified as foci per nucleus for each dose. c) Focus distribution analyzed 30 min, 1 h and 2 h post irradiation (1, 2 Gy). Results were pooled and graphs were used to compare the size distribution of foci upon each treatment. d) Representative images of γH2AX foci formation (green). Slides were counterstained with DAPI (blue) to visualize nuclei. e) Representative images of blots after incubation with the respective antibodies. f) Protein levels of p-ATM, p-p53, p-DNA-PK, and RAD51 at 1 h after 2 Gy irradiation. Results were presented as relative to GAPDH. Data are expressed as means ± SD from at least three independent experiments. The significance level was p < 0.05 (*) (two-tailed Student’s t-test) for all, except (c) where a non-parametric test (Mann-Whitney test) were used. The significance level was p < 0.05 (*), p < 0.01 (**) and p < 0.001 (****).

## 3. Results

### 3.1. Cytotoxic effects of nicotine on BEAS-2B cells

Nicotine cytotoxicity was assessed by two schemes of treatment; 1 h treatment with a dose range of 0–400 μM nicotine (Fig. 1A), and 16 h treatment with a dose range of 0–8 μM nicotine (Fig. 1B) (Lee et al., 2018; Ma et al., 2014). Consistent with previous reports, no significant toxicity was observed within the tested dose range in BEAS-2B cells (Sun et al., 2022). Considering the subsequent irradiation, 2 μM nicotine treatment starting 16 h beforeirradiation was determined asthe optimal condition to minimize cellular stress.

### 3.2. Dose-response effect of alpha-irradiation on BEAS-2B cells

BEAS-2B cells were subjected to alpha irradiation at doses of 0, 0.5, 1, 2, and 4 Gy. Fig. 1C illustrates the obtained dose-response survival curve. 1 and 2 Gy of alpha irradiation were identified as the optimal doses for this study, to achieve a cellular response in this relatively radioresistant cell line. To study the inflammatory response, additional doses of 0.5 and 4 Gy of alpha particles were also employed.

### 3.3. Nicotine aids in the preservation of cell survival in cells exposed to alpha particles

At 24 h, a decrease in viable cell number was observed in both the alpha-irradiated and the combination group compared to control (Fig. 1D). However, the combination group exhibited a slightly faster recovery trend in cell growth starting from day 4 compared to the alpha-irradiated group. This trend persisted throughout the 10-day observation period (Fig. 1F). Enhanced cell growth was associated with consistently higher average viability in the combination group compared to the alpha-irradiated group, however these differences were not statistically significant (Fig. 1E, G).

To further investigate the impact of nicotine on alpha-irradiated cells, clonogenic survival assay was conducted using doses of 1 and 2 Gy (Fig. 1H, I). Survival fraction was lower in the alpha-irradiated group (0.89 for 1 Gy and 0.52 for 2 Gy) as compared to the combination group (1.15 for 1 Gy and 1.02 for 2 Gy), highlighting the effect of nicotine in preserving cell survival in irradiated cells. This difference was statistically significant at 2 Gy (p-value = 0.0038). These data align the cell growth and viability results and suggest a possible survival-enhancing effect of nicotine on cells exposed to alpha particles.

To investigate modulatory effects of hormones on this interaction, BEAS-2B cells were treated with 10 nM DHT or E2 after irradiation. A slight increase in cell viability was observed upon treatment with DHT in all groups. Although E2 treatment showed a minimal increase as well, in nicotine alone and alpha-irradiated groups, the elevation was less than after DHT (Figure S1A). Clonogenic survival showed a similar trend (Figure S1B). Therefore, hormonal treatment appeared to not have a significant impact on cell survival or viability in response to alpha irradiation or the combination of alpha and nicotine.

### 3.4. Nicotine reduces the number and size of γH2AX foci induced by alpha particles

We then investigated any possible involvement of nicotine in processing the alpha particle-induced lesions by assessing the kinetics of γH2AX foci formation and decay. Fig. 2 illustrates the net focus frequency per cell as a function of time over a period of 24 h after 1 Gy (A) and 2 Gy (B) of alpha irradiation. The total radiation-induced foci (total foci, TF) were subdivided into small foci (SF) and large foci (LF). The TF distribution showed a biphasic γH2AX response with peaks at 1 and 6 h in the alpha-irradiated group, whereas the combination group seemed to present reduced number of foci at the time of alpha peaks, notably at 1 h (and significantly after 1 Gy). This reduction was observed in both SF and LF although it was not statistically significant. The nicotine-treated group did not show any notable induction of foci formation.

Next, we compared the size distribution of SF and LF in the alpha-irradiated and the combination groups (1 Gy, 2 Gy) in the initial hours of processing DNA damage when most of the foci have emerged by using 30 min, 1 h and 2 h focus area raw values. A difference is noticeable in the range of LF size in 1 Gy alpha-irradiated cells (ranging from 60 to 443 pixels, median =92) compared with the combination group (ranging from 60 to 331 pixels, median =85), and in 2 Gy alpha-irradiated cells (ranging from 60 to 458 pixels, median =87) compared with the combination group (ranging from 60 to 259, median =83). These data indicate that nicotine may alter the size of foci in alpha-irradiated cells by shifting them to smaller size (Fig. 2C).

### 3.5. Nicotine diminishes the persistence of damage signaling

To assess if the effect of nicotine on alpha-induced lesions extends throughout the entire repair pathway, the levels of phosphorylated DDR proteins including DNA-dependent protein kinase, catalytic subunit (DNA-PKcs) (pSer2056), ataxia telangiectasia mutated (ATM) (pSer1981), tumor protein p53 (pSer15) and radiation sensitive protein 51 (RAD51) were analyzed at 1 h post exposure to 2 Gy alpha irradiation (Fig. 2F). Although, ATM and p53 remained activated upon irradiation in both alpha-irradiated and combination groups, the latter group exhibited significant decreases in phosphorylation of the respective proteins. Similarly, phosphorylation of DNA-PKcs was reduced in the combination group compared to alpha-irradiated cells, although not statistically significant. A similar decrease was also observed in the expression level of Rad51 (p-value =0.11), however, unlike p-ATM, p-p53, and p-DNA-PKcs, the levels of RAD51 did not seem to be elevated upon irradiation in neither group at this early time point.

### 3.6. Less DNA fragmentation in alpha-irradiated cells treated with nicotine

To explore whether the decrease in repair protein levels in the combination group is associated with higher levels of unrepaired DNA, comet assay was employed to quantify the DNA fragmentation levels. DNA fragmentation was explored at the earliest time possible post exposure (3 min) up to 180 min. Although the background tail intensity was not optimal in the control group (14.25), yet, already at 3 min post irradiation the percentage of DNA present in the comet tail, tail intensity (TI), ofthe combination group(11.15) wasless thanthe alpha-irradiated group (16.79), showing less fragmented DNA (Fig. 3B, S2A). The same trend can be seen at all tested time points, up to 180 minutes, however only significant at 3, 60 and 120 min post irradiation. These data challenged our previous hypothesis of possibility of higher levels of unrepaired DNA. Instead, it leaves us with two alternative scenarios: either the damage induction was lower from the beginning, or the repair process occurred much more rapidly than anticipated after 2 Gy of alpha radiation, although then likely with increased error rates. Considering the total radiation exposure duration of 8 minutes and 58 seconds, it is possible that the formation of foci may have emerged during the irradiation period, or for nicotine, already during 16 h treatment, as previously suggested by others (Ginzkey et al., 2012; Sollazzo et al., 2017), following whichthe repair has initiated. Figure S2A shows the same data of TI, represented by the mean value of each experiment.

**Fig. 3.**
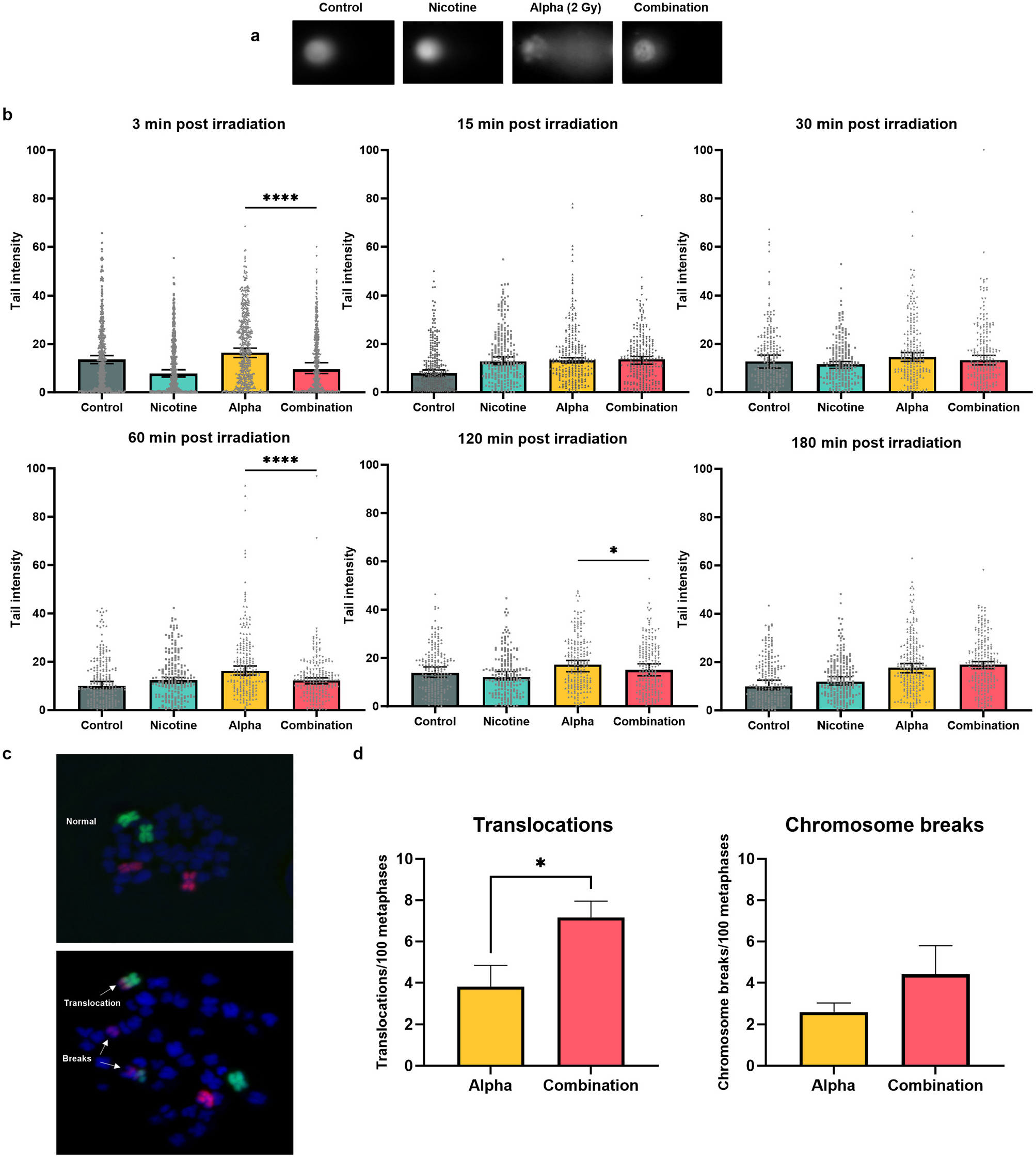
Nicotine allows faster repair while increasing the risk of translocations in alpha-irradiated cells. a) Representative images of comet heads and tails in each treatment group. b) Tail intensity (TI) upon treatment with 2 μM nicotine (pre- and post-treatment), alpha radiation (2 Gy) and combination using the alkaline comet assay. Each dot corresponds to the TI of one cell. c) Representative images of translocations and chromosome breaks. d) Frequency of translocations and chromosome breaks involving chromosome 1 (orange) and chromosome 2 (green) with each other or with other chromosomes stained with DAPI (blue), analyzed 27 h post irradiation (2 Gy). Data are expressed as means ± SD except for TI where median values are represented. Data are from at least three independent experiments. The significance level was p < 0.05 (*) and p < 0.001 (****). Statistical analysis was performed with non-parametric, Mann-Whitney test in the case of comet assay, and two-tailed Student’s t-test in the case of FISH assay.

### 3.7. Effect of nicotine on erroneous repair of alpha-induced DNA lesions

To examine if nicotine treatment can place the damaged cells at greater risk of error-prone repair, we evaluated the frequency of translocations in both the alpha-irradiated group and the combination group using FISH assay. The frequencies of chromosomal aberrations have consistently served as a reliable indicators of DNA damage (Cornforth and Loucas, 2019; Braselmann et al., 2005). In particular, we focused on stable-type chromosomal aberrations, as they have long-term implications for cancer induction. We analyzed the frequency of translocations involving chromosome 1, harboring approximately 8 % of the human entire genome (Gregory et al., 2006), and chromosome 2, as the second largest human chromosome (O’Connell et al., 1989), with each other or with other chromosomes stained with DAPI. The results in Fig. 3D shows that the combination group exhibited a higher percentage of translocations compared to the alpha-irradiated group (p-value =0.0107). Moreover, there was a slight increase in the number of chromosome breaks in the combination group. These data can suggest a role of nicotine in increasing the likelihood of erroneous repair when interacting with alpha-induced DNA lesions.

Studies have shown that alpha irradiation can cause G2/M arrest (Guerra Liberal et al., 2022). In addition, nicotine has been shown to eliminate cell cycle arrest caused by DNA damage (Lee et al., 2005). To ensure that the difference in translocations is not due to a possible S or G2 arrest post irradiation in the alpha-irradiated group and to verify the reproducibility of these findings at later time points, we conducted a preliminary test at 36 h post irradiation. Consistent results were observed with our previous findings (one replicate, Figure S3) showing increased translocation rate in the combination group.

### 3.8. Nicotine does not affect the entry into G2 arrest at 24 h post radiation but assists in recovering from the arrest at 48 h post radiation

To further investigate whether the lower translocation frequencies observed in the alpha-irradiated group can be attributed to G2 arrest (with fewer cells proceeding to metaphase) cell cycle distribution was analyzed at 4, 24 and 48 h post irradiation. At 4 h, no noticeable alterations were observed between any groups. An increase in the percentage of cells in the S/G2 phase was observed after 24 h in both alpha-irradiated and combination groups, although not statistically significant (Fig. 4A, B). This finding most likely rules out the possibility of differences in cell cycle distribution as a confounding factor in the analysis of translocations at 27 h (24 h plus 3 h colcemid treatment). However, a slight decrease in the percentage of cells in S/G2 was observed after 48 h in the combination group compared to the alpha-irradiated group (Fig. 4C). This indicates an eliminating effect of nicotine on the cells in G2 arrest after 48 h post irradiation, and may serve as an initiator of the faster recovery as validated by cell growth and viability data (Fig. 1E, F, G). Irrespective of nicotine pretreatment, alpha exposure (1 Gy) also triggers G1 arrest as judged from accumulation of the p53-responsive transcript CDKN1A at 24 h (Fig. 4D), likely falling between the 4 and 24 h cell cycle analysis time points.

**Fig. 4.**
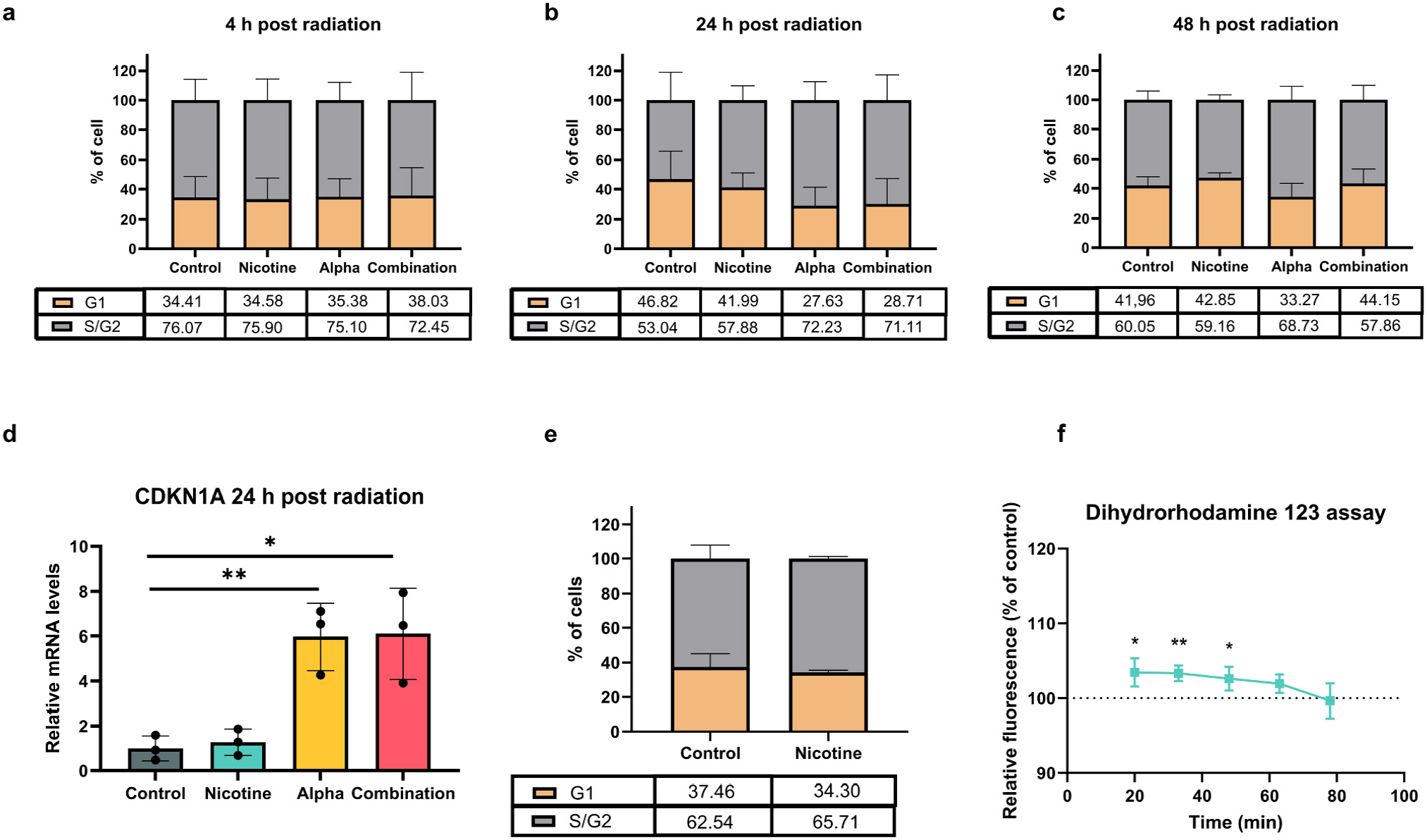
Nicotine does not affect the entry into S/G2 arrest but assists in recovery from it. a-c) Cell cycle distribution at 4 h (a), 24 h (b) and 48 h (c) post irradiation (2 Gy) in cells treated with 2 μM nicotine (pre- and post-treatment), alpha irradiation, or the combination. d) mRNA levels of CDKN1A at 24 h post alpha radiation. e) Cell cycle distribution upon 16 h treatment with 2 μM nicotine. f) RNOS/ROS generation after 16 h treatment with 2 μM nicotine measured after 20 minutes of incubation and every 10 minutes afterwards in presence of dihydrorhodamine 123. Data are expressed as means ± SD from at least three independent experiments. The significance level was p < 0.05 (*) and p < 0.01 (**) (two-tailed Student’s t-test).

Furthermore, to address the hypothesis that cells could be in a less radiosensitive cell cycle phase at the time of exposure, we assessed if cell cycle alterations were present at the time of exposure, after 16 h nicotine pre-treatment. No significant difference was observed between the control and nicotine-treated group, demonstrating that 16 h nicotine pre-treatment did not affect the status of cell cycle at the time of irradiation (Fig. 4E).

### 3.9. Nicotine induces a very modest increase in mitochondrial production of ROS

Previous studies have demonstrated that nicotine can influence the initiation of ROS production (Ramalingam et al., 2021; Yogeswaran and Rahman, 2022) and oxidative stress might be involved in the induction of DNA damage through oxidation of DNA bases (Wu et al., 2005). To further explore the role of nicotine at the doses employed in this study, we analyzed the level of RNOS/ROS induction in BEAS-2B cells after 16 h of 2 μM nicotine pre-treatment using the DHR123 assay. BEAS-2B cells treated with nicotine exhibited a 3 % increase in RNOS/ROS formation compared to the control group (Fig. 4F).

### 3.10. Inflammation can be another mechanism through which nicotine and alpha-particles interact to promote lung cancer

Given that inflammation contributes to cancer development and supports various stages of tumorigenesis (Greten and Grivennikov, 2019), it becomes essential to explore the possible interplay between nicotine and alpha particles in affecting inflammatory cytokines. We investigated alterations in mRNA levels of two pro-inflammatory cytokines, IL-6 and IL-1β, previously shown to be implicated in the progression and/or initiation of lung cancer (Qu et al., 2015; Zhang and Veeramachaneni, 2022).

As shown in Fig. 5A, mRNA levels of IL-6 began to rise at 3 h and were elevated until 6 h post irradiation in both alpha-irradiated and combination groups. At 24 h after exposure, the levels of this proinflammatory marker had returned to baseline levels. Conversely, mRNA levels of IL-1β started to increase at 6 h and remained elevated at 24 h post irradiation, indicating a delayed yet sustained response compared to IL-6. Considering the clear dose-response trend observed at 3 h post radiation for IL-6 and 6 h post radiation for IL-1β, these time points were selected as critical for drawing conclusions. While both the alpha-irradiated and combination groups exhibited an up-regulation of IL-6 and IL-1β genes, the combination group demonstrated higher levels of these pro-inflammatory markers, particularly at higher doses (4 Gy). Although the data was not statistically significant, the increased trend can still suggest a supportive effect for nicotine in inducing proinflammatory cytokines when interacting with alpha particle-induced damage.

**Fig. 5.**
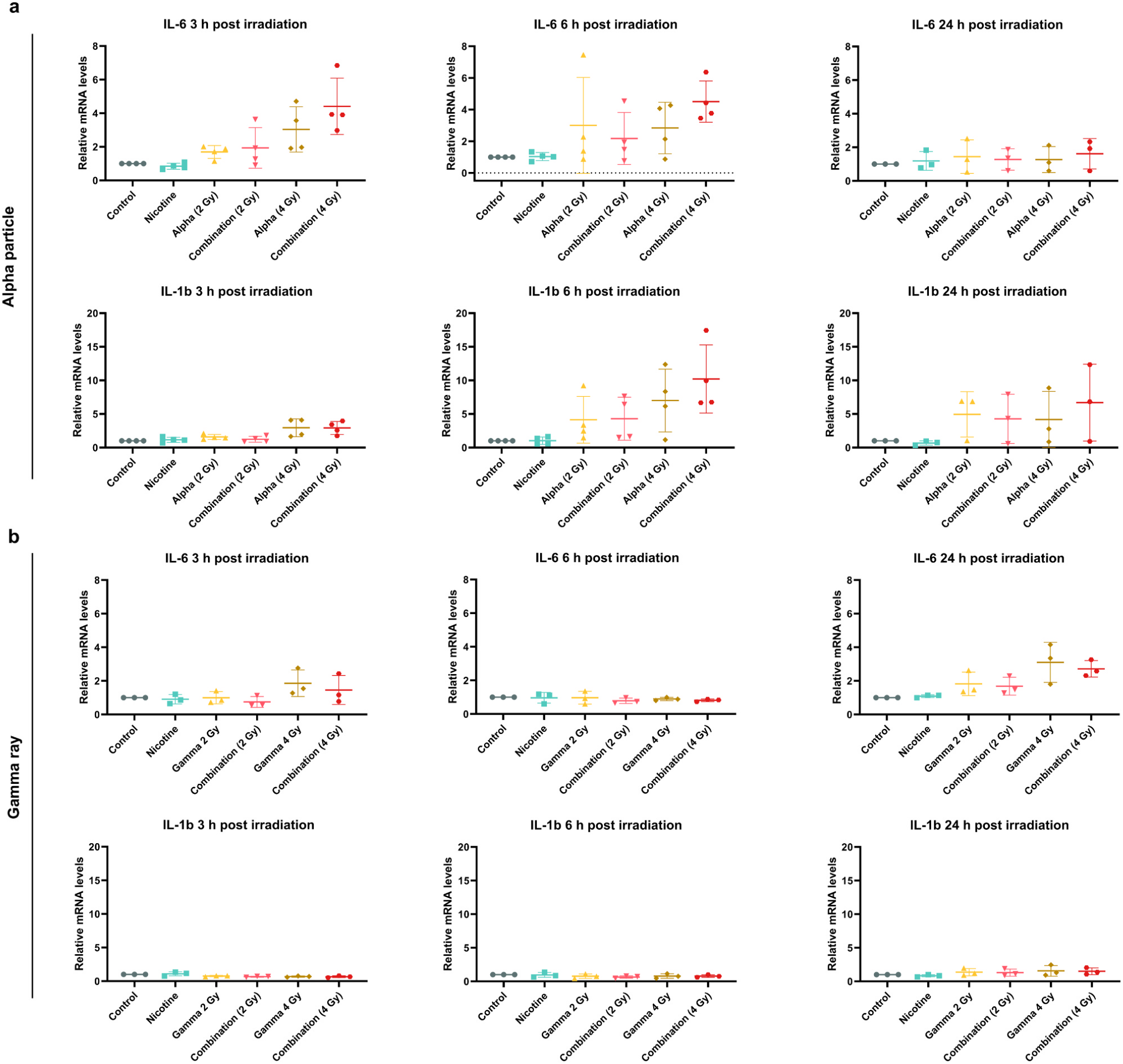
mRNA levels of pro-inflammatory genes. IL-6 and IL-1β mRNA levels were analyzed in BEAS-2B cells 3, 6 and 24 h post irradiation with 2 and 4 Gy of alpha particles (a) or gamma rays (b), 2 μM nicotine (pre- and post-treatment), or the combination. Data are expressed as means ± SD from at least three independent experiments.

### 3.11. Nicotine demonstrates distinct effects in response to different radiation qualities

To compare the interaction of nicotine with high and low-LET irradiation, mRNA levels of aforementioned pro-inflammatory cytokines were assessed after gamma irradiation. While there was a slight, late increase in IL-6 levels in both groups 24 h post irradiation, there was no noticeable induction of IL-6 or IL-1β observed in either group at the other studied time points, suggesting that radiation response and nicotine interaction may differ between low and high LET radiation (Fig. 5B). To further explore this, γH2AX foci formation and decay kinetics were assessed (Fig. 6A, B), and compared with alpha particle irradiation data (Fig. 2A, B). Interestingly, there were no significant differences in TF between the gamma-irradiated and combination groups after 1 Gy. However, the combination group showed a delayed reduction trend in the number of LF, starting at 2 hours post irradiation, with less noticeable changes in SF as compared with alpha-irradiated cells. With 2 Gy gamma irradiation, there was instead a slight initial elevation observed, mainly in SF, in the combination group during the 1–2 hours post-irradiation period.

**Fig. 6.**
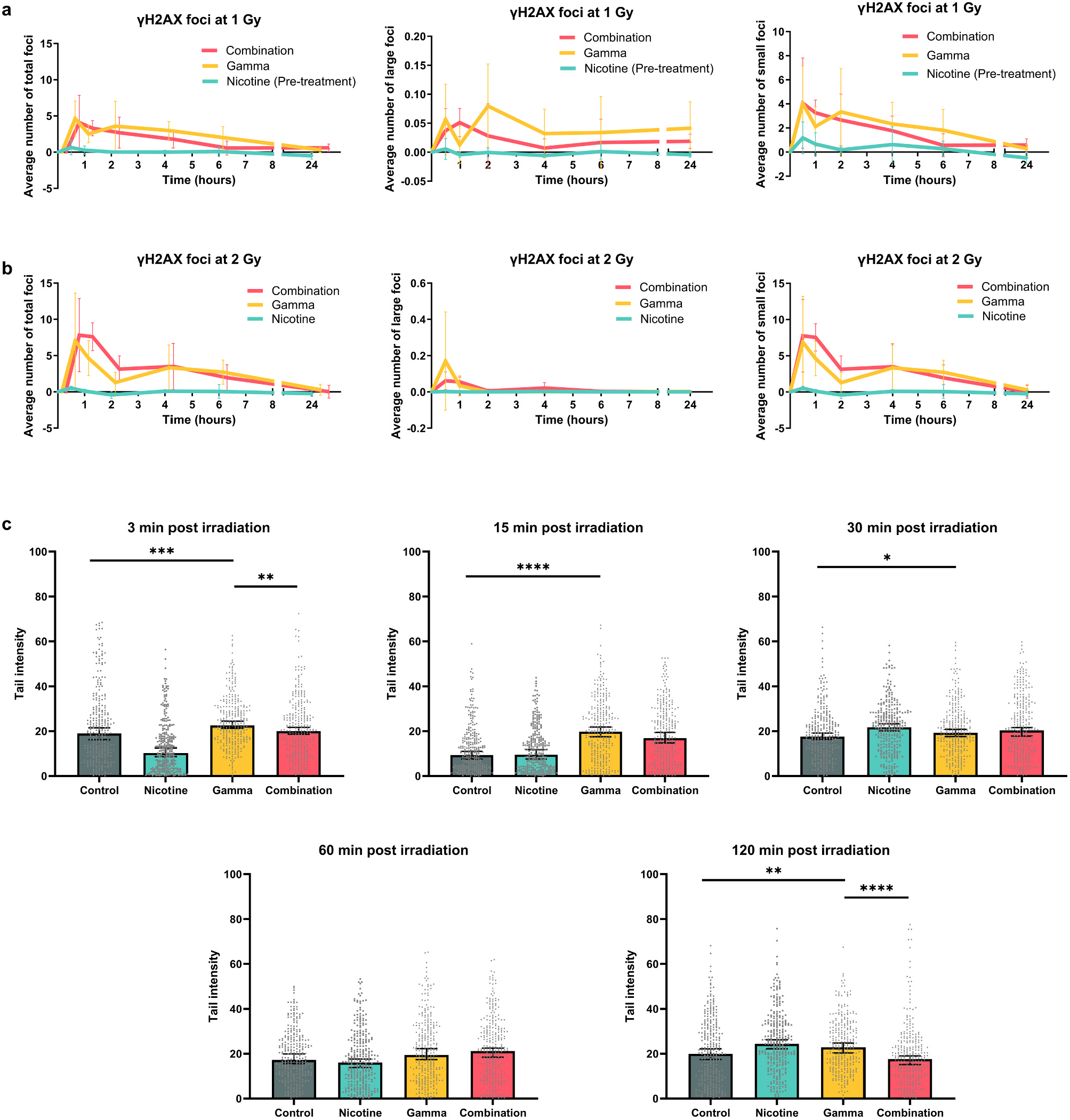
Distinct effects of nicotine in response to gamma irradiation. a, b) Kinetic response of γH2AX foci formation as a function of time over 24 h. Immunofluorescence was used to capture representative γH2AX focus images from control, gamma-irradiated (1, 2 Gy), and 2 μM nicotine-treated cells (pre-treatment for 1 Gy, pre- and post-treatment for 2 Gy). γH2AX foci were quantified as foci per nucleus for each dose. c) Tail intensity (TI) upon treatment with nicotine (pre- and post-treatment), gamma radiation (2 Gy) and combination using the alkaline comet assay. Each dot corresponds to the TI of one cell. Data are expressed as means ± SD except for TI where median values are represented. Data are from at least three independent experiments. A non-parametric test (Mann-Whitney test) were used to analyze the comet assay data. The significance level was p < 0.05 (*), p < 0.01 (**), p < 0.005 (***) and p < 0.001 (****).

The unique interplay between nicotine and different radiation qualities was further supported by comet assay data upon exposure to gamma radiation. As depicted in Fig. 6C, there was a minor decline in TI during the initial 15 minutes post irradiation in the combination group compared tothe gamma-irradiated group. This declining trendcould not be observed again until the final time point at 120 minutes post irradiation, where a significant decrease was noticeable. The nicotine interaction with foci induced by gamma rays exhibited a delayed impact of nicotine on reducing TI, highlighting minimal to no difference between the groups at earlier time points in contrast to the alpha-irradiated data (Fig. 3B).

### 3.12. Effect of chromatin modifications on nicotine interaction with alpha particles

To determine the possible mechanisms by which nicotine interacts with alpha-induced damage, heterochromatin alterations were assessed. Since effective DNA repair encounters obstacles in the form of heterochromatin (Goodarzi and Jeggo, 2012), we aimed to explore whether our observed DDR signaling alteration in the combination group could result from a different chromatin structure, as nicotine is suggested to support chromatin alterations consistent with permissive state by increasing histone acetylation and decreasing histone methylation (Muenstermann and Clemens, 2024). To that end, first histone acetylation and methylation were assessed by measuring the protein levels of H3K9ac, H4K8ac and H3K9me3 at 1 h post irradiation. Fig. 7A shows a slight reduction in the level of H3K9me3 as well as an insignificant increase in the H4K8ac level in the combination group in comparison with the alpha-irradiated group, suggesting a trend towards supporting histone acetylation and against methylation. However, H3K9ac levels do not seem to follow the same trend. To further explore this, PU139 was utilized to investigate whether nicotine retains its effects when acetylation is suppressed by evaluation the levels of two crucial DDR proteins, p-p53 and γH2AX. Fig. 7C validates the diminished acetylation following a 16 h treatment with 5 μM PU139, confirming the efficacy of the drug. As Fig. 7E depicts, in the PU139 treated groups, both p-p53 and γH2AX protein levels were not following the same reduction trend in the combination group as compared with alpha-irradiated group, when acetylation is inhibited. These data further imply the possible role of histone acetylation as a contributing factor in the interaction between nicotine and DNA damage induced by alpha particles.

**Fig. 7.**
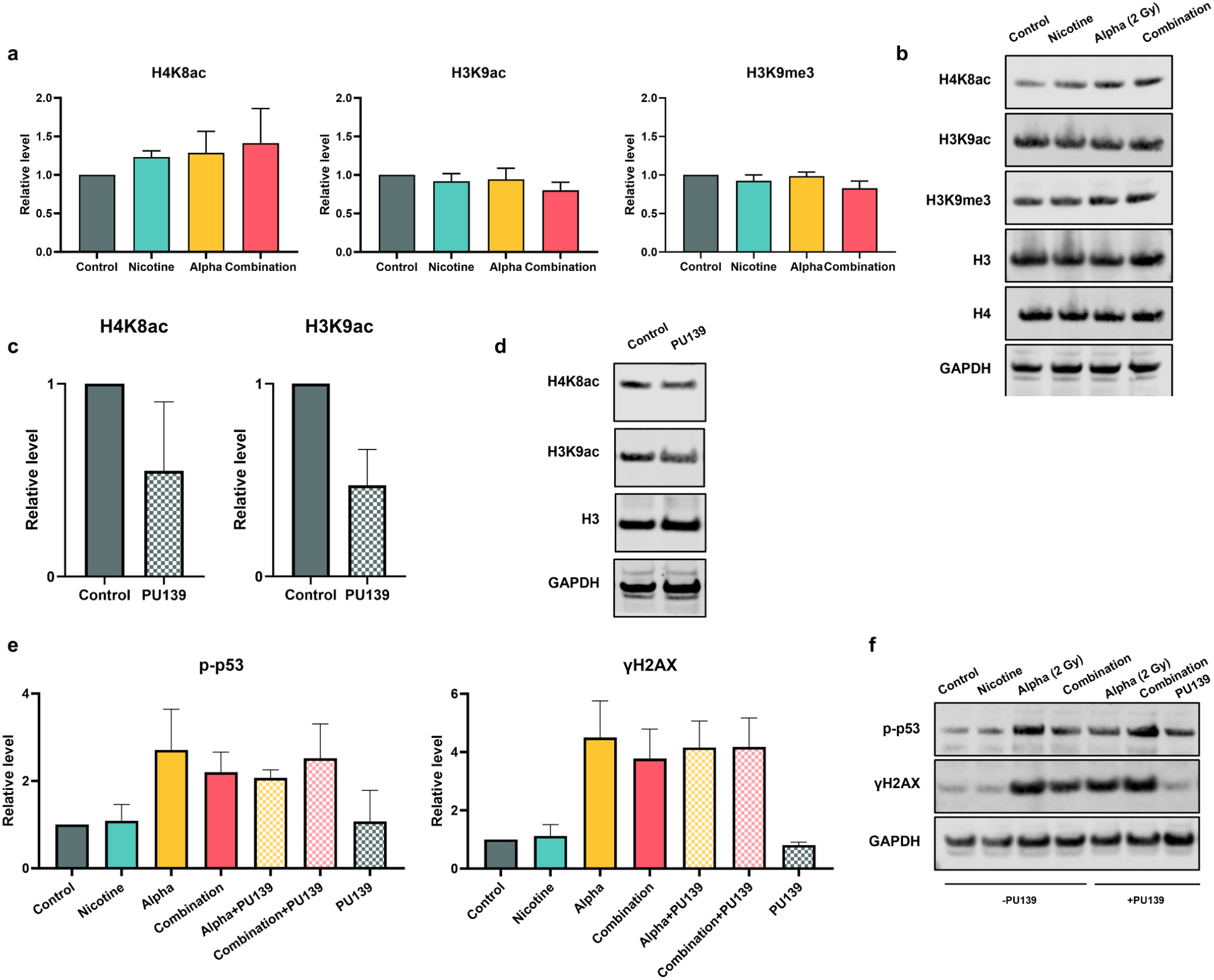
Chromatin modification involvement in nicotine interaction with alpha particles. a) Protein levels of H4K8ac, H3K9ac and H3K9me3 at 1 h after 2 Gy alpha irradiation. b, d, f,) Representative images of blots after incubation with the respective antibody. c) Protein levels of H4K8ac and H3K9ac following treatment with 5 μM PU139 for 16 h. e) Protein levels of p-p53 and γH2AX in different groups of treatments with or without PU139 1 h after alpha irradiation (2 Gy). Results were presented as relative to GAPDH, and H4K8ac, H3K9ac, H3K9me3 were further normalized to either H4 or H3. Data are expressed as means ± SD from at least three independent experiments.

## 4. Discussion

Epidemiological studies on miners as well as residents of areas with high radon levels have been the principal sources of evidence on the combined effect of CS and radiation (Darby et al., 2005; Leuraud et al., 2011). However, limited numbers of studies have been defining underlying mechanisms involved in this interaction. Although the carcinogenic potential of vaping or using nicotine products may be orders of magnitude lower than CS, the popularity of these “safer” alternatives has attracted research attention to embark on the carcinogenic effect of them (Goniewicz et al., 2014). This study was designed as an *in vitro* experiment to assess the molecular interaction between nicotine and different radiation qualities by investigating its impact on processing the DNA lesions and other systemic mechanisms in favor of cancer promotion including oxidative stress and inflammation. We reasoned that focusing on nicotine would benefit a large population of users at risk of lung cancer development. Our study provides new evidence to suggest a role of nicotine in rapid recovery of BEAS-2B cells exposed to alpha irradiation while increasing the likelihood of erroneous repair.

Evidence from a number of studies indicate that nicotine enhances the tumor promoting properties of cells by induction of proliferation and invasion in a variety of human cancer cell lines, including lung cancer (Dasgupta et al., 2009). In addition, nicotine has been shown to support lung tumorigenesis through perturbing cellular surveillance (Zhang et al., 2020). However, in normal BEAS-2B cells, our study indicated that nicotine alone, at the doses and schedule utilized, did not exert a notable impact on the mechanisms under investigation, except a slight growth promotion when supplied continuously. Nevertheless, when cells were exposed to alpha particles, nicotine produced an accelerated recovery trend and increased cell survival. In line with our data, nicotine promoted growth and tumorigenesis caused by DNA-damaging agents such as benzopyrene and γ-irradiation by enhancing cyclin D1 expression and attenuating Chk2 activation, leading to reduced G1 arrest (Nishioka et al., 2011). In accordance with this finding, a dose response behavior was seen for radon or CS alone, and greater in the combination, in enhancing the colony forming capacity of BEAS-2B cells (Huang et al., 2017).

Chromatin is highly dynamic in proliferating cells due to replication and mitosis, but also transcription (Ma et al., 2015). It is generally accepted that histone acetylation is correlated with decompacted chromatin, and serves as an active mark for gene activation by allowing RNA polymerases to pass through (Henikoff and Shilatifard, 2011). H4K8ac is associated with stimulation of transcription initiation (Gupta et al., 2017) and is considered a major regulator of chromatin-linked transcriptional alterations (Radzisheuskaya et al., 2021). Furthermore, acetylation of histone H4 residues K8 and K12 has been shown to have a pivotal role in loosening chromatin structures during the process of DNA replication (Ruan et al., 2015). Given the marginally improved cell growth and recovery observed in the combination group, the slight elevation in H4K8ac levels may support the presence of a transcriptionally active state and a more relaxed chromatin structure which can be further supported by the reduced level of H3K9me3, a marker of peripheral heterochromatin (Towbin et al., 2012). Despite being a marker of active promoter state (Karmodiya et al., 2012) and active transcription (Xu et al., 2022), H3K9ac does not follow this trend.

DNA repair processes are conditional on chromatin landscape and accessibility (Mansisidor and Risca, 2022). A permissible chromatin state allows prompt opening of chromatin in the vicinity of the lesions and efficient recruitment of repair proteins and initiation of the repair process (Cleaver, 1977; Murr et al., 2006). The MRN complex (Mre11, Rad50, Nbs1), is crucial at the site of damage for the phosphorylation and subsequent activation of ataxia telangiectasia mutated (ATM) (Uziel et al., 2003), which phosphorylates the Ser139 site on H2AX proteins. In this study, both ATM, H2AX and p53 were activated in the groups irradiated with alpha particles, however, the combination group exhibited significantly less phosphorylation. These data together with trends towards reduction of the levels of p-DNA-PK for the NHEJ pathway, and RAD51 for the HR pathway in the combination group, indicate an attenuated DDR response at 1 h post exposure.

The complex interplay between nicotine, DNA damage and repair mechanisms has been explored in other studies. However, interpretations regarding the impact on less fragmented DNA or suppression of repair proteins have varied. Studies have suggested that nicotine and its metabolites impair the repair system by reducing the levels and activities of DNA repair proteins XPC and OGG1/2 after higher doses in BEAS-2B and UROtsa cells (via proteosomal/autophagosomal degradation), as well as in lung tissues derived from mice (Lee et al., 2018). However, it remains unclear from our study whether this decrease in repair protein activation results from an instant response that has already been resolved by the time of assessment, or stems from a suppression of the repair process (wherein more fragmented DNA should persist). Nicotine inhibited DNA strand breaks induced by genotoxic nitrosamine NNK in the hepatic cell line HepaRG (Ordonez et al., 2014). In our study, the level of DNA fragmentation, quantified by comet assay, showed a diminished TI in the combination group already 3 min post irradiation which was in support of the first hypothesis, suggesting that nicotine might have initiated the repair process already during the irradiation time. This data was further supported by the influence of nicotine on the size of foci induced by alpha particles, causing a shift towards smaller foci sizes in the irradiated cells. However, the expected higher number of LF in the 2 Gy alpha group compared to the 1 Gy group was not detected. A possible reason might be the limitations due to the resolution of fluorescent microscopy. When numerous LF are in close proximity, the emitted light may overlap, causing challenges for the software in distinguishing individual foci.

The lower activation of ATM, p53 and p-DNA-pk in the combination group may therefore reflect that the peak came earlier than after 1 h, and this would be more in line with the radioresistance by clonogenic survival. Cell cycle differences were likely not strong contributors for differences seen by γH2AX or western blot analysis at 1 h since there was no change at either 0 or 4 h. Although nicotine treatment only led to a slight (3 %), but significant increase in RNOS/ROS production after 16 h, it is possible that the oxidative stress was elevated earlier. An increased tail moment was present at 1 h after 1 mM nicotine in BEAS-2B cells, and a similar induction was inhibited using an antioxidant in nasal mucosa cells (Ginzkey et al., 2012). ROS can lead to the formation of bulky lesions by causing base oxidation and DNA breaks, that can stall the replication fork (Branzei and Foiani, 2010; Sancar et al., 2004). Defects in DNA replication can lead to chromosome rearrangements by affecting the replication fork stability and progression, thereby influencing replication-associated repair processes, and increasing the risk of chromosomal aberrations (Branzei and Foiani, 2010; Bruhn and Foiani, 2019). Although ROS-mediated mechanisms could be another possible pathway through which nicotine could contribute to the development of chromosomal aberrations, the slight increase in ROS upon nicotine treatment might be a supportive, but not primary, mechanism by which nicotine influences DNA damage or its response.

To our knowledge, no study exists on nicotine interaction with alpha particle-induced DNA lesions. Nevertheless, a study by Lee et al., have reported an additive behavior of alpha particles and CS in lung epithelial cells (including BEAS-2B) in induction of γH2AX foci (Lee and Kim, 2019). However, such results may not be directly translatable to our study due to different study setup including assessment of the whole CS extract, not only nicotine, as well as difference in the dose ranges of alpha particles (up to 0.25 Gy, sourced from ^241^Am, with dose rates of 0.079 Gy/min, while, our studies employed doses of 1 and 2 Gy, administered at a dose rate of 0.223 Gy/min).

An accelerated onset of repair associated with a permissible state of chromatin in the combination group can explain the less fragmented DNA and reduction of DDR protein activation. Interestingly, following acetylation inhibition, we could not observe the same effect of nicotine in reducing damage signaling in the combination group, shown by assessing the protein levels of p-p53 and γH2AX as two important DDR proteins. This result emphasizes the possibility that the observed nicotine effects might have been associated with chromatin state. Given the data supporting a permissible chromatin state together with rapid repair, an open chromatin state can possibly contribute to an increased risk of chromosomal translocations. Translocations are persistent aberrations that are known as a hallmark of cancer (Hogenbirk et al., 2016). DNA surrounding the DSBs needs to be accessible to the other broken end of double-stranded DNA for translocations to occur (Hromas et al., 2016), making the translocation breakpoints enriched with open chromatin marks (Roukos and Misteli, 2014). Accordingly, the open chromatin state in the combination group can further explain the possibility of a higher translocation rate. In addition, it has been shown that monomethylation of histone 3 lysine 4 (H3K4me1) can give rise to translocations in multiple models, while the trimethylation form of histone 3 lysine 9 (H3K9me3) can reduce the rate of chromosomal translocations (Burman et al., 2015; Makova and Hardison, 2015), which is aligned with our results showing a slight reduction in H3K9me3 levels in the combination group with higher rate of translocations. Coherent with our study, investigations on the effect of nicotine on chromosomal aberrations have shown that nicotine leads to higher levels of chromosomal aberrations in human fetal cells in vitro using amniotic cells grown in nicotine containing medium (25 ng/ml) (Demirhan et al., 2011). These data can support a role of nicotine in increasing the likelihood of erroneous repair when interacting with alpha-induced DNA lesions.

Nevertheless, cells are equipped with cell cycle checkpoints to prevent cells from proliferating with genomic instabilities, by causing cell cycle arrest (Kastan and Bartek, 2004). Factors that override the cell cycle arrest can give rise to tumor progression by securing a sustained proliferation of damaged cells (Kastan and Bartek, 2004), and, it has been widely held that nicotine can eliminate cell cycle arrest caused by DNA damage (Lee et al., 2005; Nishioka et al., 2011). Along similar lines, in our study, although nicotine treatment did not affect the entry into G2 arrest, a faster recovery from G2 arrest caused by irradiation was detected at 48 h post irradiation in the combination group as compared with the alpha-irradiated group. This data was consistent with cell proliferation data showing faster growth and recovery in the combination group. Higher translocation rates together with shortened G2 arrest and higher proliferative and survival potential observed in the combination group elucidate a tumor-supportive behavior of nicotine when interacting with alpha-induced DNA lesions.

Furthermore, the comparison between the interaction of nicotine with different radiation qualities showed that nicotine could not notably alter the formation of γH2AX foci, TI and the level of pro-inflammatory markers when interacting with gamma ray-induced damages. Given the distinct behavior of nicotine in response to different radiation qualities together with the negligible alterations observed subsequent to nicotine treatment alone, it appears that nicotine mostly influences cellular responses to DNA damage when a higher proportion of DSBs is present. This notion arises from the limited impact observed in the gamma radiation data where a larger proportion of the DNA damage is indirect, forming through reactive species from water hydrolysis. In fact, the radiation doses employed for gamma radiation were equivalent to those used for alpha particles in terms of absorbed dose, regardless of the higher relative biological effectiveness of high-LET radiation, leading to less DNA damage potential in case of gamma irradiation. In support of this, a divergent response with a faster repair after alpha radiation compared to gamma radiation was noted in breast cancer cells when an open chromatin was induced by HDAC inhibition (Svetlicic et al., 2020). Nevertheless, conducting more comprehensive studies is essential to further validate our findings and shed a light on the interaction between nicotine and chromatin modifications.

In addition, our data showed a trend towards up-regulation of IL-6 and IL-1β genes in a dose-response manner in the combination group. In high-LET scenarios, achieving statistically significant results is challenging due to diverse and non-homogeneous particle behaviors. Consequently, high error bars are common, likely even more for rapid alterations like immune responses. This led us to also consider consistent trends in our discussion. Although studies have claimed a dual effect of nicotine with both pro- and anti-inflammatory properties (Zhang et al., 2022), coherent with our results, increased inflammation and cell invasion was observed in oral squamous cell carcinoma cells exposed to electronic cigarette vapor extract (Robin et al., 2022). Multiple studies have reported that ROS and pro-inflammatory markers, such as IL-6, can contribute to chemotherapy resistance through various mechanisms including up-regulation of anti-apoptotic genes, stimulation of stemness properties, and enhancement of DNA repair processes (Liu et al., 2016; Chen and Chang, 2019; Savant et al., 2018). Interestingly, in line with these findings, our data on pro-inflammatory markers accompanied by the data of increased oxidative stress, less fragmented DNA, higher rate of translocations and possibly accelerated repair processes in the combination group can help to clarify the interplay between nicotine and DNA lesions induced by radiation, which can have implications for accurately estimating radiation risk in the general population, including nicotine users. Given the observed interaction between nicotine and alpha particles, it would be highly desirable to mechanistically further explore the involvement of error-prone DSB repair pathways in translocation formation and the possibility of involvement of alternative repair pathways (which may pose a higher risk of erroneous repair) due to the suppression of common repair pathways (cNHEJ, HR). Also, conducting similar end points in other respiratory cell lines with different concentrations of nicotine and doses of alpha irradiation is recommended, if applicable.

## 5. Conclusions

In conclusion, the results of the study indicate that nicotine interacts with DNA lesions induced by alpha particles by inducing a faster but more error-prone repair as well as shortened cell cycle arrest, which may increase the risk of occurrence of translocations and secure the proliferation of cells with genomic instabilities. One possible mechanism by which nicotine may exert its function is through chromatin modifications that favor a more relaxed chromatin structure. This hypothesis is supported by the fact that inhibiting histone acetylation compromised nicotine ability to function similarly as it did previously in processing the DNA lesions induced by alpha particles. Other systemic alterations can also support nicotine in exerting its effects including oxidative stress and inflammation. Overall, our study points towards the necessity of considering nicotine when transferring lung cancer risks among the general population for radiation exposure.

## CRediT authorship contribution statement

**Nadia Boroumand:** Writing – original draft, Visualization, Methodology, Investigation, Formal analysis, Data curation. **Lovisa Lundholm:** Writing – review & editing, Supervision, Project administration, Funding acquisition, Conceptualization. **Karine Elihn:** Writing – review & editing, Supervision, Conceptualization. **Carol Baghdissar:** Investigation, Formal analysis.

## Data availability

Data will be made available on request.

## Acknowledgements

We are grateful to Andrzej Wojcik from the Department of Molecular Biosciences, The Wenner-Gren Institute, Stockholm University, for his kind contribution and input in designing and performing the study, as well as helpful discussions concerning the radiation field.

## Appendix A. Supporting information

Supplementary data associated with this article can be found in the online version at <u>doi:10.1016/j.ecoenv.2024.117009</u>.

